# Saracatinib synergizes with enzalutamide to downregulate androgen receptor activity in castration resistant prostate cancer

**DOI:** 10.1101/2023.04.22.537922

**Authors:** Ralph E. White, Maxwell Bannister, Abderrahman Day, Hannah E. Bergom, Victor M. Tan, Justin Hwang, Hai Dang Nguyen, Justin M. Drake

**Author notes:** **Correspondence :** Justin M. Drake.

## Abstract

Prostate cancer (PCa) remains the most diagnosed non-skin cancer amongst the American male population. Treatment for localized prostate cancer consists of androgen deprivation therapies (ADTs), which typically inhibit androgen production and the androgen receptor (AR). Though initially effective, a subset of patients will develop resistance to ADTs and the tumors will transition to castration-resistant prostate cancer (CRPC). Second generation hormonal therapies such as abiraterone acetate and enzalutamide are typically given to men with CRPC. However, these treatments are not curative and typically prolong survival only by a few months. Several resistance mechanisms contribute to this lack of efficacy such as the emergence of AR mutations, AR amplification, lineage plasticity, AR splice variants (AR-Vs) and increased kinase signaling. Having identified SRC kinase as a key tyrosine kinase enriched in CRPC patient tumors from our previous work, we evaluated whether inhibition of SRC kinase synergizes with enzalutamide or chemotherapy in several prostate cancer cell lines expressing variable AR isoforms. We observed robust synergy between the SRC kinase inhibitor, saracatinib, and enzalutamide, in the AR-FL+/AR-V+ CRPC cell lines, LNCaP95 and 22Rv1. We also observed that saracatinib significantly decreases AR Y^534^ phosphorylation, a key SRC kinase substrate residue, on AR-FL and AR-Vs, along with the AR regulome, supporting key mechanisms of synergy with enzalutamide. Lastly, we also found that the saracatinib-enzalutamide combination reduced DNA replication compared to the saracatinib-docetaxel combination, resulting in marked increased apoptosis. By elucidating this combination strategy, we provide pre-clinical data that suggests combining SRC kinase inhibitors with enzalutamide in select patients that express both AR-FL and AR-Vs.

## 1 Introduction

Prostate cancer remains the highest diagnosed non-skin cancer amongst the American male population and is second highest in deaths, next to lung cancer. When primary prostate cancer is diagnosed in patients, many clinicians will either prescribe surgery, radiotherapy, and/or androgen deprivation therapy (ADT) that interfere with androgen synthesis such as leuprolide (GnRH agonist)(1). While potentially curative in up to 70-80% of men, the remaining 20-30% of men will develop tumors that become resistant to ADT with a rising prostate specific antigen termed castration resistant prostate cancer (CRPC). Newer, second generation therapies such as enzalutamide (AR competitive antagonist)(2, 3) and abiraterone acetate (CYP17A inhibitor)(4) have been developed to prolong survival in both hormone-naïve and hormone-resistant cancer. However, none of these agents are curative, prompting investigators to elucidate the major mechanisms of resistance in CRPC.

Previous literature has implicated several major mechanisms of resistance in CRPC such as the amplification of the androgen receptor (AR), loss of *PTEN*, and *TMPRSS-ERG* fusions (5–7). One key mechanism is the emergence of androgen receptor splice variants (AR-Vs) (8). AR-Vs are constitutively-active truncated versions of AR that lack the C-terminal ligand binding domain and can function independently in the presence of androgen, leading to additional AR transcriptional activity. Clinically, AR-Vs (in particular AR-V7) increase in abundance and are implicated in resistance to prior hormonal therapies such as abiraterone acetate and enzalutamide (9). Recent studies show chemotherapy, such as docetaxel, has a higher treatment efficacy in patient tumors expressing AR-V7(10). Yet, these therapies are toxic, leading to side effects that heavily impact quality of life for the patient. Chemotherapy also does not inhibit primary mechanisms of resistance involving AR, revealing the need for alternative approaches to treatment.

Another mechanism of resistance is increased tyrosine kinase signaling (11). Our previous work evaluated the phosphoproteome of CRPC patients at autopsy. Using multi-omic integration, we took a kinase-centric approach to identify SRC kinase as a key activated kinase and signaling hub in CRPC (12). SRC kinase, also known as c-SRC, is a non-receptor tyrosine kinase that plays major roles in cancer cell proliferation, communication, and adhesion (13, 14). Specifically to prostate cancer, SRC kinase phosphorylates AR at Y^534^ (15) and directly interacts with the AR N-Terminal domain via hydrophobic interactions with SRC’s SH3 domain (16). Phosphorylation of AR by SRC kinase maintains AR stability and transcriptional activity (17, 18) along with regulating other kinases that phosphorylate other residues of AR via growth factor stimulation and kinase crosstalk (19–21). While intriguing as a pre-clinical target, SRC kinase inhibition in clinical trials for treatment of CRPC has not been successful (22). Administration of a dual SRC kinase and BCR-Abl inhibitor, dasatinib, failed in late stage CRPC clinical trials as both a monotherapy and in combination with docetaxel (23). These clinical trial results dampened the excitement around SRC kinase as a viable target in CRPC. However, several explanations may exist as to why these trials were not successful, such as lack of patient stratification and broadly targeted combinations that do not focus on AR inhibition.

To resolve this, in this study, we provide pre-clinical data that supports SRC kinase inhibition with standard of care hormonal therapies such as enzalutamide for treating AR positive CRPC. We find that enzalutamide plus saracatinib was strongly synergistic in AR-full length positive (AR-FL+) cell lines, regardless of AR-V positive (AR-V+) status. Meanwhile, docetaxel plus saracatinib was not as effective, especially in cell lines expressing AR-Vs, supporting the potential failure of dasatinib in the previously mentioned clinical trial with docetaxel. We also found that saracatinib ablated AR Y^534^ phosphorylation, AR-V protein expression, and altered AR specific gene signatures, suggesting that AR stability and transcriptional activity was perturbed through SRC kinase inhibition. Lastly, we also observed that saracatinib induced higher levels of γH2AX, DNA replication stress when in combination with enzalutamide, and markers of apoptosis in an AR-FL+/AR-V+ cell line 22Rv1.

## 2 Results

### 2.1 Enzalutamide and saracatinib yields strong synergy in AR-FL+ cell lines

To begin studying our drug combinations, we selected prostate cancer cell lines that expressed SRC kinase with different AR genetic backgrounds (Figure 1A), expecting a myriad of responses that will allow us to evaluate synergy between our selected drugs. Using these cell lines, we generated dose-response curves to determine the IC50s of enzalutamide (enza), docetaxel (DTX), and the SRC kinase inhibitor saracatinib (sara) (Figure 1B). Using the IC50 dosage, we administered serial dilutions of enza plus sara or DTX plus sara to our cell lines (Supplemental Figure 1A). Synergy was then calculated via Bliss Independence (BI) and the Combination Index (CI) equation via Chou Talalay method through CompuSyn 1.1 (24, 25). AR-FL+ only AD1 cells showed synergy in both enza plus sara and DTX plus sara combinations (Figure 1D, Supplemental Figure 1B). Synergy was also observed in both AR-FL+ and AR-V+ 22Rv1 and LNCaP95 cells (Figure 1E, F, Supplemental Figure 1B) with the enza plus sara combination. Interestingly, synergy was observed in LNCaP95 cells with the DTX plus sara combination but not in 22Rv1 cells. AR-V+ R1D567 cells showed synergy in the enza plus sara combination via the BI model (Figure 1F) and showed additivity via the CI model (Supplemental Figure 1B). There is a lack of synergy seen between DTX plus sara in R1D567 using both models (Figure 1G, Supplemental Figure 1B). Using CompuSyn 1.1, we were able to generate a dose reduction index (DRI), which determines the fold reduction of each drug when in combination with another drug. In our 22Rv1 and LNCaP95 cell lines, we see a reduction up to 5-fold for both enza and sara when in combination (Supplemental Figure 1C). We also observe a 3-fold reduction for DTX and a near zero fold reduction for sara when in combination, which suggests that the reduction of sara was more potent in the presence of enza versus DTX. Lastly, we show no synergy, but rather antagonism, between enza plus sara and DTX plus sara in the AR negative DU145 cells (Figure 1H). We expected no synergy for enza plus sara in DU145 cells due to enza’s inability to enact its mechanism of action because of DU145’s lack of AR. Overall, these findings indicate there is substantial synergy between enza and sara in PCa cell lines and this synergy potentially coincides with the presence of AR-FL alone or in the presence of AR-V expression.

**Figure 1:**
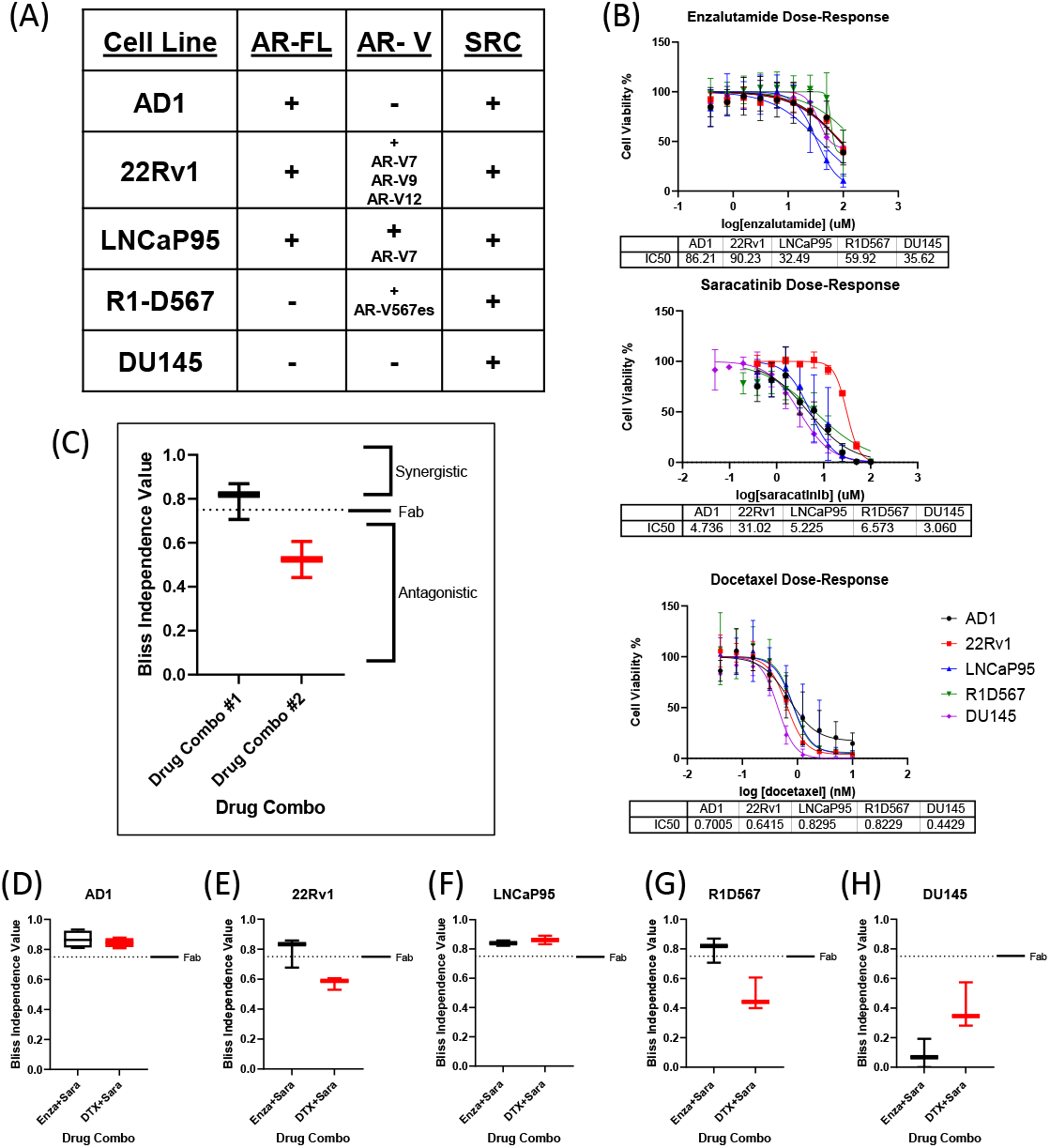
Synergy observed between Enzalutamide and SRC kinase Inhibitors in AR+ Positive Cell Lines. (A) Cell lines used for in vitro studies and corresponding AR status and SRC status. (B)Dose-response curves for Enzalutamide (Enza), Saracatinib (Sara), and Docetaxel (DTX) for each cell line. IC50 dosage of each drug to each cell line used is shown below each curve (N≥ 3). (C) Representative Bliss Independence graph for drug combinations. Bliss Independence values > than Fab are considered synergistic (D-G) Bliss independence results for drug combos for cell lines (AD1, 22Rv1, LNCaP95, R1D567, DU145). N≥3 for all cell lines except LNCaP95 where N=2.

### 2.2 Saracatinib decreases AR phosphorylation and AR-V protein expression via SRC kinase inhibition

Previous literature has shown that certain kinases can regulate AR function, stability, and activity via phosphorylation on particular residues of AR. For example, CDK1, 5, and 9 phosphorylates AR S^81^, which is located within the N-terminal domain of AR and regulates AR transactivation, transcription, and nuclear localization (26–28). SRC kinase phosphorylates AR on residue Y^534^, which is critical for AR stability and transcription (17, 18). To assess the role of SRC kinase inhibition on AR phosphorylation and how that may contribute to drug synergy in CRPC, we seeded the CRPC cell lines 22Rv1 and LNCaP95 overnight followed by a media change to charcoal stripped serum media for 3 days. We then administered individual and drug combinations for 24 hours at each drug’s IC50 followed by stimulation with R1881, a synthetic androgen, and epidermal growth factor (EGF) for 5 minutes to induce maximal phosphorylation on AR. We found that sara ablated the phosphorylation of AR Y^534^ (both AR-FL and AR-Vs) in both 22Rv1 and LNCaP-95 cells (Figure 2A, B). This effect was sara dependent as enzalutamide and docetaxel were unable to decrease this phosphorylation site, while sara alone and in combination with enza or DTX all produced similar reduction of this phosphorylation residue. We also measured AR S^81^ phosphorylation and found that its reduction coincided with the reduction of AR protein. Interestingly, we also found that total AR-V protein expression was decreased by sara, so we measured AR-V7 and found decreased expression in sara-treated samples. Overall, these findings indicate that pharmacological ablation of SRC kinase can heavily affect AR phosphorylation and AR protein expression.

**Figure 2:**
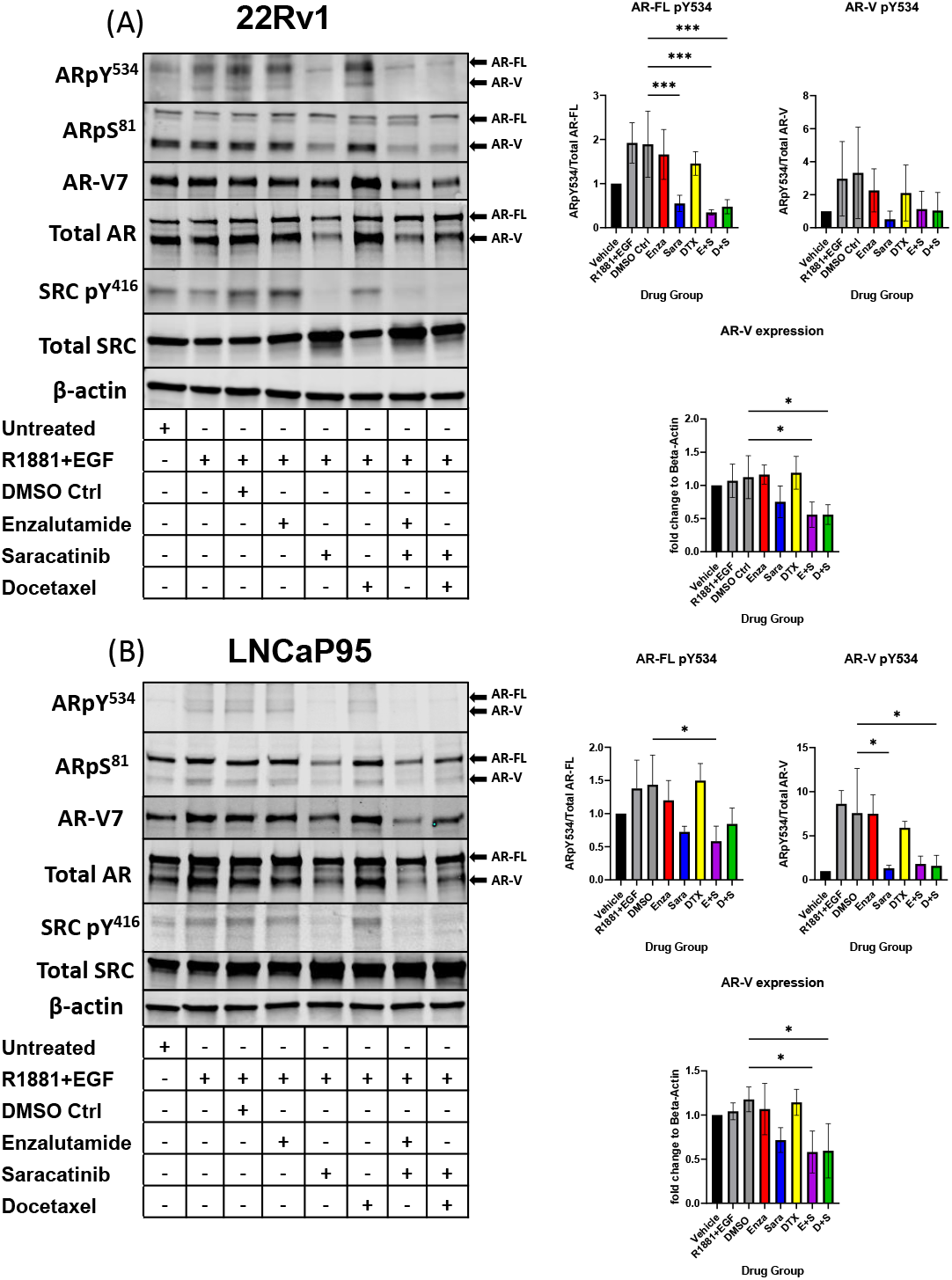
AR Y^534^ phosphorylation and AR-V protein expression ablated via SRC kinase inhibition. (A) 22Rv1 and (B) LNCaP95 cells seeded in FBS media for 24hrs then switched to Charcoal-Stripped Serum (5%) RPMI Media and grown over 3 days. Cells are then given the drugs for 24hrs and stimulated with R1881, a synthetic androgen, and Epidermal growth factor (EGF) for 5 minutes. Drug groups are DMSO, Enza, Sara, DTX, Enza plus Sara (E+S), and DTX plus Sara (D+S). Blotting for AR, SRC, ARpY534, ARpS81, SRCpY416, β-actin (N≥ 3) and ARv7 (N=2). *, P < 0.05; **, P < 0.01; ***, P < 0.001; and ****, P < 0.0001.

### 2.3 Saracatinib alters AR gene signature in CRPC

Based on previous literature that stated phosphorylation on AR Y^534^ regulated AR transcriptional activity, we decided to perform RNA-seq to evaluate the consequence of AR-specific gene signatures after sara-induced reduction of AR Y^534^. 22Rv1 cells were prepared as stated previously and administered drug groups (enza, sara, enza plus sara) were added. Cells were harvested after 48 hours and RNA sequencing was performed. We initially focused on changes in steroid receptor mRNA expression as it had been reported that inhibition of AR can induce expression of another steroid receptor as a mechanism of resistance to AR targeted therapies, such as with the glucocorticoid receptor (NR3C1) and mineralocorticoid receptor (NR3C2) (29, 30) We also utilized gene signatures from select literature. We found that sara reduced AR mRNA expression (Figure 3A) and enza induced the expression of the glucocorticoid (NR3C1) and mineralocorticoid (NR3C2) receptors, similar to what was published previously. We also found that the enza plus sara combination further reduced AR mRNA expression with little induction of NR3C1 and NR3C2 expression, hinting towards sara preventing this mechanism of resistance. We also observed that sara altered AR gene signature activity in both the Dehm (31) and Nelson (32) AR-regulated gene sets. In the Dehm gene set, which contains 19 AR-regulated genes, we found that sara treatment alone reduced certain AR-regulated genes (e.g. ABCC4, KLK3, ACSL3, and ELL2) and the combination of sara and enza dramatically reduced the expression of several AR-regulated genes that were less perturbed by either treatment alone, including ZBTB10, PMEPA1, CENPN, NKX3-1, and FKBP5 (Figure 3B). Similar effects were also observed in the Nelson gene set with sara-dependent reduction identified unique AR-regulated genes when compared to the sara plus enza combination. We also found that sara can reduce the AR-regulated gene signature scores in both the Dehm and Nelson gene sets similar to enza and that the combination of enza plus sara even further reduced the AR activity signatures scores in both datasets (Figure 3B, C). Overall, these findings indicate that SRC kinase ablation can heavily affect AR dependent gene expression and may work with enzalutamide to further inhibit AR-specific activity.

**Figure 3:**
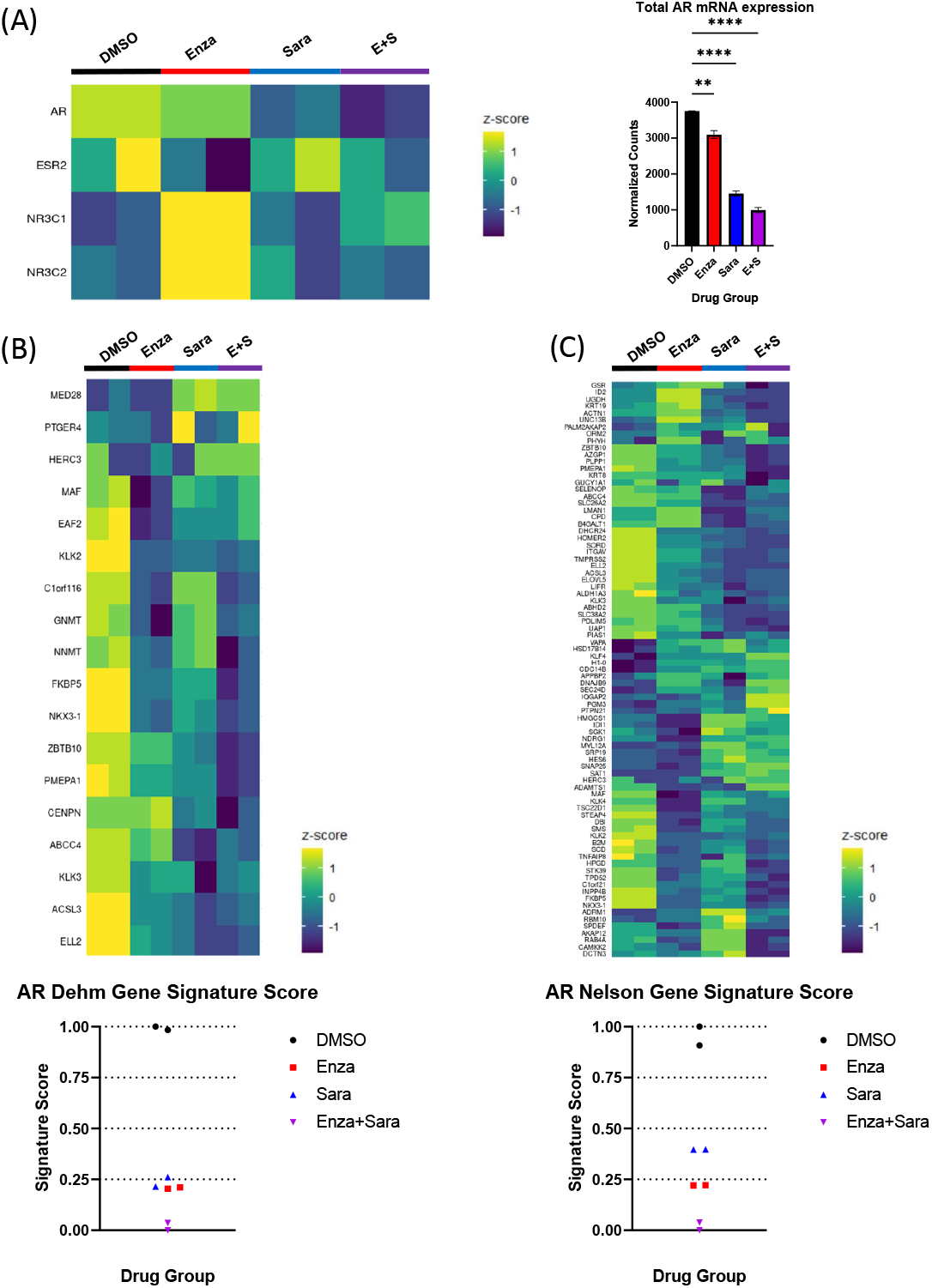
Saracatinib affects AR gene signatures. 22Rv1 seeded in FBS media for 24 hrs then switched to Charcoal-Stripped Serum (5%) RPMI Media and grown over 3 days. Cells are then given the drugs, R1881, and EGF for 48hrs. Drug groups are DMSO, Enza, Sara, or E+S. (A) Heatmap showing steroid receptor gene expression (left) with corresponding bar graph showing normalized counts of AR mRNA expression (right). (B) Heatmap of Dehm gene signature (19 genes) corresponding to AR activity with signature score (C) Heatmap of Nelson AR gene signature(83 genes) with signature score. N=2 *, P < 0.05; **, P < 0.01; ***, P < 0.001; and ****, P < 0.0001.

### 2.4 Saracatinib induces DNA damage and apoptosis via DNA replication stress

Since sara alone and in combination with enza significantly lowered AR activity signature scores, and that blocking AR function can induce cell death via apoptosis and suppression of cell growth (33–35), we sought to investigate if sara plus enza synergized in exerting their cytotoxic effects during the cell cycle and activating apoptotic pathways. First, to follow the cell cycle status of individual cells in asynchronous growing cell populations, we stained the chromatin-bound PCNA, a component of the DNA replication fork, and DNA in the 22Rv1 cell line using antibody and DAPI, respectively (Supplemental Figure 2A, left, Figure 4A). We also pulse-labeled newly synthesized DNA with EdU, a modified thymidine nucleoside incorporated into the DNA of actively proliferating cells. Cells undergoing DNA replication displayed high levels of chromatin-bound PCNA and became EdU-positive after pulse-labeling. We then established a cell cycle profile using quantitative image-based cytometry, comparing PCNA and DAPI in each drug group to identify G1, S, and G2 cell populations (Supplemental Figure 2A, left). Sara treatment alone or in combination with enza or DTX, caused a modest, but statistically significant, decrease in S phase cells (Supplemental Figure 2A, right). We then evaluated our drug combinations’ impact on DNA synthesis by measuring pulse-labeled EdU in PCNA-positive cells. In S phase cells (PCNA-positive), enza and sara individually decreased EdU incorporation in comparison to DTX (Figure 4B). We also saw enza plus sara decreases EdU incorporation to baseline in comparison to DTX plus sara, showing enza plus sara can greatly halt DNA synthesis. With DNA synthesis impacted, this may lead to DNA damage in S phase. We measured H2AX phosphorylation on residue S^139^, also known as γH2AX, a known marker for DNA damage. We found that the sara alone and in combination with enza or DTX significantly induced higher levels of γH2AX in S phase cells (Figure 4C). Also, γH2AX expression was not specific to any one cell cycle population, as sara alone and in combination with enza or DTX caused greater amounts of γH2AX in G1 cells (Supplemental Figure 2B, left), while enza plus sara induced the highest amounts of γH2AX in G2 cells (Supplemental Figure 2B, right). Lastly, to understand activation of apoptotic pathways, we measured activated caspase 3/7 and cleaved PARP, early markers for apoptosis required for late stages in apoptosis. We detected higher levels of activated caspase 3/7 in the enza plus sara combination over each drug alone (Figure 4D) as well as higher cleaved PARP expression (Figure 4E). Overall, these findings indicate SRC kinase inhibition via sara causes DNA replication stress that is supplemented more by enzalutamide vs. docetaxel, resulting in greater activation of apoptotic pathways.

**Figure 4:**
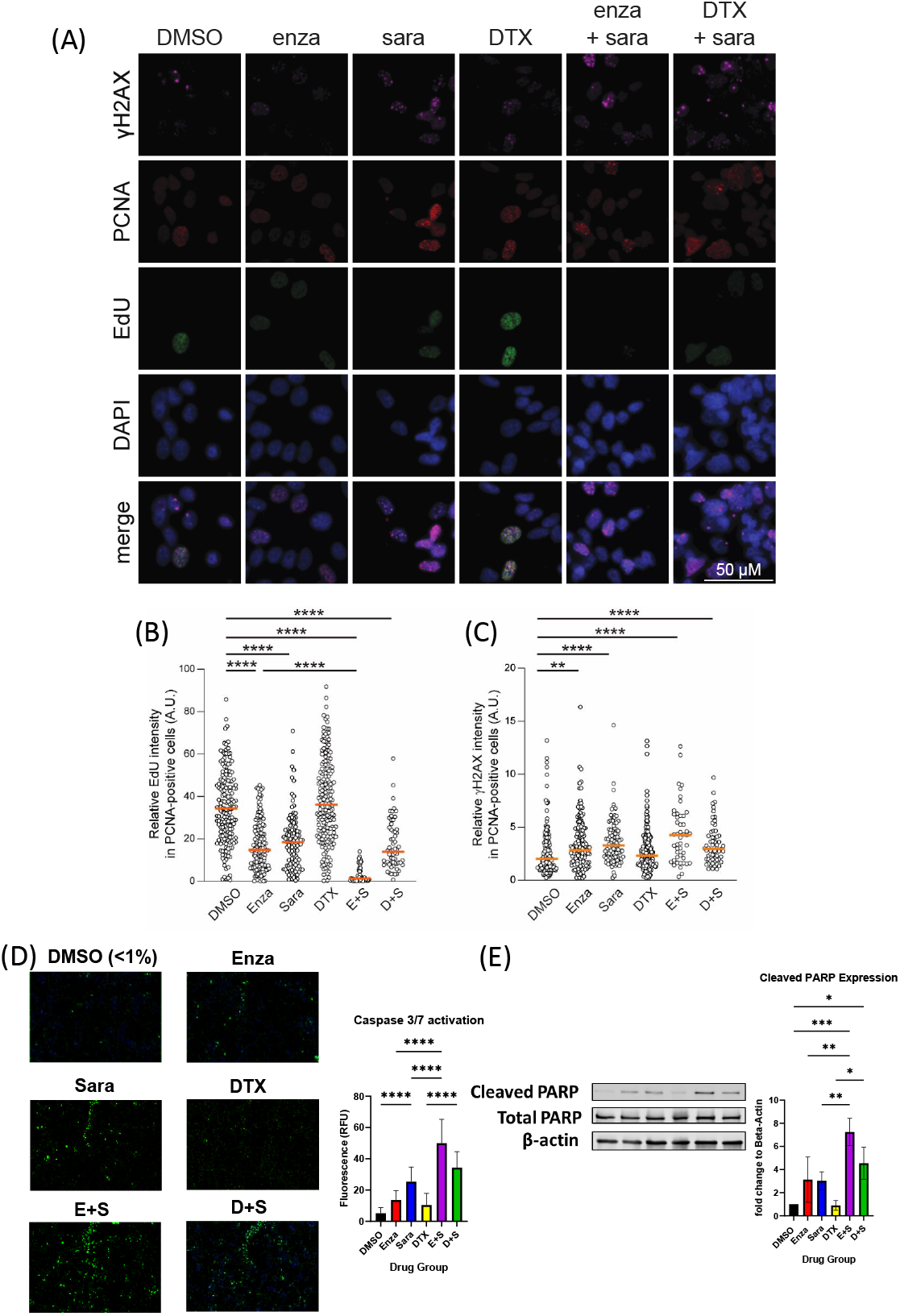
Saracatinib halts DNA synthesis, induces DNA Damage and activates markers of apoptosis. (A) Representative images of charcoal-stripped 22Rv1 cells treated with are DMSO, Enza, Sara, DTX, E+S, or D+S, for 24 hrs then stained for indicated antibodies. (B) EdU (via Click-It) and (C) γH2AX quantification. (D) caspase 3/7 activation (via Cell Event) representative, detected via immunofluorescence. (E) Western blot of Cleaved PARP, Total PARP, and β-actin. N=3. *, P < 0.05; **, P < 0.01; ***, P < 0.001; and ****, P < 0.0001.

## 3 Discussion

In this study, we found that enzalutamide plus saracatinib is synergistic in prostate cancer cell lines that express AR-FL, regardless of AR-V status. We also found that docetaxel plus saracatinib is synergistic to additive in cell lines that express AR-FL only. However, synergy is reduced between docetaxel and saracatinib in cell lines that express AR-Vs, highlighting its treatment potential in the correct context. Overall, we exemplify how AR status can affect the efficacy of combination therapy *in vitro*. Clinically, this is important for the patient as we can identify the appropriate combination strategy to have a greater impact on halting their disease progression and circumvent any unnecessary side effects. Under castrate conditions, AR can shift from its canonical ligand dependent signaling to non-canonical ligand independent signaling (15, 36, 37). This adaptive signaling can lead to the expression of constitutively active AR-Vs and increased tyrosine kinase activity. SRC kinase phosphorylates AR and has been shown to bind to the N-Terminal domain of AR, leading to a gain in cell proliferation and signaling (15). We found that saracatinib decreased AR Y^534^ on both AR-FL and AR-Vs and, interestingly, saracatinib also decreased AR-V protein expression, including AR-V7. This points out that changes in phosphorylation on AR Y^534^ may affect AR-V protein stability highlighting a possible route towards its ligand-independent function.

Previous literature has shown that the phosphorylation of AR can activate and contribute to AR-dependent gene networks (26–28, 37). In particular, Y^534^ phosphorylation by SRC kinase contributes to AR transcriptional activity and nuclear localization (21). Therefore, we evaluated saracatinib’s effects on AR mRNA expression alongside other steroid receptors’ mRNA expression. We found that AR mRNA expression decreased in the presence of saracatinib and an increase in GR and MR mRNA expression when enzalutamide was given. The effect of enzalutamide is blunted when combined with saracatinib, suggesting that saracatinib affects AR-gene regulated specific mechanisms of resistance tied to enzalutamide. We also observed that saracatinib altered AR gene signature scores to a similar level as enzalutamide, which was quite surprising. We found many of the major AR target genes were impacted, such as TMPRSS2, KLK2/3, and FKBP5. While this mechanism requires more investigation, we postulate that saracatinib impacts AR-V protein stability, resulting in reduced AR-V binding to DNA that in return lessens activation of ligand-independent AR gene expression.

AR inhibition has been shown cause DNA damage on telomeres to prostate cancer cells in previous literature(33, 35), so we evaluated our drug combinations in DNA synthesis, DNA damage, and apoptosis. Using quantitative immunofluorescence cytometry, we identified G1, S phase, and G2 cell populations in each of our drug groups and found that saracatinib decreased the percentage of cells in S-phase. We also observed that enzalutamide and saracatinib, individually and together, halted EdU incorporation greater than docetaxel alone, indicating that cells undergo DNA replication stress when given this drug combination. We also found that saracatinib induced greater γH2AX expression vs. enzalutamide or docetaxel. Lastly, we found greater caspase 3/7 activity and cleaved PARP expression in cells administered enzalutamide and saracatinib.

While the *in vitro* mechanisms combining sara plus enza are quite striking and point towards SRC kinase as a key therapeutic target in CRPC, clinical trial data has been not as positive. In the phase 3 clinical trial, READY, docetaxel plus the dual SRC kinase and BCR-Abl inhibitor, dasatinib, were given to metastatic CRPC patients who were naïve to chemotherapy (23). While the trial was unable to meet the specified primary endpoints and no benefit in overall survival, it should be noted that there was a lack of CRPC patient stratification by AR or SRC kinase status. It should also be pointed out that while docetaxel is a standard of care in CRPC (9), it does not directly affect AR function or activity. From a clinical perspective and our work presented here, the combination of enza plus sara would benefit a portion of CRPC patients who still retain AR activity, including AR-Vs, over docetaxel plus sara. Our study may also prompt investigators to look deeper into select kinases that phosphorylate AR, such as CDK1/5/9, ACK1, SRC, and MAPK (21), as inhibition of these kinases in AR+ prostate cancers may provide patient benefit in combination with AR-targeted agents.

Evaluating combination therapy, in comparison to monotherapy, by using mathematical equations, such as Bliss Independence and Combination Index, can possibly bridge the clinical gap for testing (38). Both equations note trends of synergy and the interplay of the drugs’ mechanisms of actions based on the cell model (24, 25). Specifically, for our purposes, we focused on AR species as the variable of our models and saw different responses with enzalutamide and docetaxel when paired with saracatinib. While enzalutamide combinations were synergistic and resulted in greater prostate cancer cell death, it is still important to note that docetaxel combinations were synergistic to additive in some of our models, citing the importance of using docetaxel in the clinic in some cases, especially when AR-Vs are not expressed.

Major topics of interest to build upon these findings involve the accessibility of AR and deciphering the mechanism of DNA damage caused by saracatinib. Since saracatinib alters AR transcriptional activity, SRC kinase inhibition could cause chromatin remodeling of AR as well as affect co-activator recruitment, as shown in previous literature with CDKs (27). Saracatinib’s induction of γH2AX is of key importance to determine if saracatinib causes DNA replication forks and how that impacts DNA repair pathways. Overall, we present SRC kinase inhibition as a therapeutic strategy to be combined with current AR therapies available for use to treat AR driven CRPC. Though we show this promising pharmacological intervention, there are limitations. With SRC kinase inhibitors, like many kinase inhibitors, they are promiscuous which lead to off target effects. This prompts a need to develop better kinase inhibitors with specificity to the chosen kinase target. Also, while saracatinib is a potent SRC kinase inhibitor, it also inhibits other members of the SRC family kinases, including Lyn and Fgr, which are also expressed in prostatic tissue. This requires further investigation into each kinases’ activity and how their inhibition could impact prostate cancer cell death.

## 4 Methods

### 4.1 Cell Culture

Human prostate cancer cell lines DU-145, 22Rv1, LNCaP95 were obtained from ATCC and cultured according to ATCC protocol in RPMI1640, supplemented with 10% FBS and 1% penicillin-streptomycin and 1% Glutamax, a substitute for L-glutamine (Corning). AD1 and R1D567 Cells were obtained from Dr. Scott Dehm at the University of Minnesota Medical School and cultured as described previously (39). Cells were not used beyond 25 passages. All cells were grown and maintained in a humidified incubator at 37°C and 5% CO2

### 4.2 Drug Dose Response (Cell Viability)

Cells were seeded at the following densities: DU-145 (500 cells/0.1 mL), AD1 (2,000 cells/0.1 mL), R1D567 (1,000 cells/0.1 mL), LNCaP95 (4,000 cells/0.1 mL), 22Rv1 (2,000 cells/0.1 mL) in 96 well plates in RPMI media (Sigma-Aldrich) with 10% FBS, 1% penicillin-streptomycin, and 1% Glutamax (Corning). After an overnight incubation at 37°C and 5% CO2, media in the wells was replaced with fresh RPMI media with 5% charcoal-stripped (CSS)-FBS, 1% penicillin-streptomycin, and 1% Glutamax (Corning). Cells were then grown in the media for 3 days. One of the following three drug groups is then administered: enzalutamide, saracatinib, ranging from 0.39-100μM, and docetaxel 0.04-10nM. All drugs were obtained from Selleckchem. Treatment lasted for 6 days with replenishment of the media and drug after 3 days. Cell viability was measured using WST-1 at 1:10 dilution with CSS-FBS media at absorbance of 450 nm (Tecan 1100 Plate Reader). IC50 dosage was calculated using GraphPad Prism. Each data point was conducted in technical and biological triplicate.

### 4.3 Synergy Studies

Cells were seeded at the densities grown as stated above. Drug combinations used included enzalutamide plus saracatinib and docetaxel plus Saracatinib at their respective IC50 dosages for each cell line. Therapy was given, beginning at 2x the IC50 dose, then serial diluted by 2 till 9 dilution groups were established. Cell viability was measured using WST-1 at 1:10 dilution with CSS-FBS media at absorbance of 450 nm (Tecan 1100 Plate Reader). Measured absorbance was converted in percentile and inputted in the Bliss Independence equation (24) as well as CompuSyn 1.1, a computer software that determines synergy via Combination Index (25) between drugs based on individual dose response. Bliss independence is calculated as Fab=Fa x Fb, where Fa/b is the fraction of cells affected by drug A/B and Fab is the product of the two fractions, representing the predicted additivity of the two drugs. Bliss independence value (represented in Figure 1C) is portrayed with Fab and the experimental values of each combination. Combinations with values greater than Fab are considered synergistic, while combination with values less than Fab are considered antagonistic. Each data point was conducted in technical triplicate.

### 4.4 Immunofluorescence Imaging and Quantitative Image-based Cytometry

For cell-cycle analysis of 22Rv1 cells, a quantitative image-based cytometry method was used as described previously (40). Briefly, cells were labeled with 10 μM EdU for 30 min and processed with the Click-IT EdU Alexa Fluor 488 Imaging Kit (Invitrogen, #C10337) according to the manufacturer’s instructions. Otherwise cells were extracted with 1x PBS containing 0.1% Triton-X100 for 10 min on ice prior to fixation with 3% paraformaldehyde/2% sucrose for 15 min at ambient temperature. Subsequently, cells were permeabilized with 100% methanol at −20°C for 10 min, blocked in blocking buffer (1x TBS containing 5% BSA, 0.05% Tween-20) for 1 hr, and incubated in primary antibodies for PCNA (mouse, 1:200, Calbiochem #PC10) and γH2AX (rabbit, 1:1000, CST #9718S) in blocking buffer overnight at 4°C. Next day, cells were washed 3 times with PBS-T before incubation with Cy5 anti-rabbit and Cy3 anti-mouse secondary antibodies for 1 hr at ambient temperature. Cells were stained with DAPI before mounting coverslips on slides.

Z-stack images were captured using a Leica DMi8 microscope (Leica Microsystems). Image segmentation of nuclei and whole cells was performed using the cellpose algorithm implemented in Python. The cyto2 and nuclei models were further trained on the images in this study to achieve high-quality segmentation. Nuclear and/or cellular masks were exported to ImageJ to measure total intensity, mean intensity, and pixel areas of defined regions within max-projected images.

### 4.5 Caspase 3/7 Activation

To measure caspase 3/7 activation, Invitrogen CellEvent Caspase-3/7 green detection reagent (C10423, ThermoFisher) was used as stated in manufacturer’s protocol. 22Rv1 cells were seeded in 96 well plates at 4,000 cells/100 μL in FBS media overnight. Media in the wells was then replaced with CSS-FBS media and grown for 3 days. The following drug groups were then administered for 24 hours at IC50 dosage: DMSO Vehicle, enzalutamide, saracatinib, docetaxel, enzalutamide plus saracatinib, docetaxel plus saracatinib. After 24 hours, the caspase reagent, diluted to 4 μM was added to the cells for 1 hour. Fluorescence was observed using Spark Cyto Multimode Plate Reader. Excitation and emission settings were 488 and 590/50 nm respectively. The intensity of fluorescence was analyzed with Spark Cyto Imaging software

### 4.6 Immunoblot Analysis

Cells were lysed with RIPA buffer supplemented with protease inhibitor tablets and phosphatase inhibitor cocktail. Protein concentration was quantified using Pierce BCA protein assay kit following manufacturer’s protocol. 40 micrograms of protein were loaded into GenScript SurePage 4%-12% polyacrylamide gel, transferred to both nitrocellulose and PVDF membranes, blocked in 5% BSA or 5% milk in 1x TBST for one hour, before incubating the membranes with primary antibodies overnight at 4°C in 1% BSA solution. Membranes were washed with 1x TBST 3 three times before incubating the membranes in LI-COR IR-conjugated secondary antibodies (1:10,000-20,000) for 2 hours at room temperature. Membranes were washed three times with 1x TBST and imaged using the LI-COR Odyssey System. Membranes were adjusted and quantified with the LI-COR Image Studio Lite software (v5.2). The following antibodies were used for Western Blot Analysis: At 1:1000 Total AR, Total SRC, Total β-actin, ARpY534, SRCpY416, cleaved PARP, total PARP. At 1:500, ARpS81, AR-V7. Blots are results in triplicate.

### 4.7 RNA sequencing and Analysis

Total RNA was isolated from 22Rv1 cells via RNeasy Mini Kit (QIAGEN, no. 74106). Messenger RNA was purified from total RNA using poly-T oligo-attached magnetic beads. After fragmentation, the first strand cDNA was synthesized using hexamer primers followed by second strand cDNA synthesis. Library sequencing was conducted on Novaseq6000 S4 flowcell for PE150 sequencing (Novogene Corporaiton Inc., Sacramernto, CA 95817). Transcriptome sequence data processing and analysis were performed using pipelines at the Minnesota Supercomputing Institute (MSI) and University of Minnesota Informatics Institute (UMII) at the University of Minnesota. Raw reads were trimmed, aligned to the GRCh38 human genome, and gene-level read counts were generated using the CHURP pipeline (41). All downstream gene expression analyses and visualizations were conducted using R (4.2.1), RStudio (2022.07.2+576) (42) and GraphPad Prism 9. Genes with less than ten total counts across all samples were filtered out. Count normalization was conducted using DESeq2’s median of ratios method (43). Visualizations were generated using the R packages ggplot2 (44) and ClassDiscovery (45), and Graphpad Prism 9.

### 4.8 Gene Activity Scoring

Relative AR activity scores were computed by summing the normalized counts of the set of genes defining each signature. Peter Nelson’s AR signature includes 83 genes, all of which were present in the dataset (29). Scott Dehm’s signature includes 19 genes, 18 of which were present in the dataset (30).

### 4.9 Statistics and Analysis

The data were presented as the mean ± SD for the indicated number of independently performed experiments, except the immunofluorescence data, which was presented as the median. The statistical significance (p<0.05) was determined using GraphPad Prism 9 with the tests indicated in the figure legends. P < 0.05 was considered to indicate a statistically significant difference. P values were determined with significance indicated as follows: *, P < 0.05; **, P < 0.01; ***, P < 0.001; and ****, P < 0.0001. Tukey’s multiple comparisons test and Kruskal-Wallis test were performed after one-way ANOVA.

## 5 Conflict of Interest

JMD has no conflicts relevant to this work. However, he holds equity in and serves as Chief Scientific Officer of Astrin Biosciences. This interest has been reviewed and managed by the University of Minnesota in accordance with its Conflict of Interest policies. None of these companies contributed to or directed any of the research reported in this article. The remaining authors declare no potential conflicts of interest.

## 6 Author Contributions

REW and JMD conceived the project. REW and VMT designed, conducted experiments and analysed data under supervision of JMD. MB performed microscopy, quantitative image-based cytometry analysis, and analysed data under the supervision of REW and HDN. HEB, AD and JHH performed RNA sequencing analysis and provided project discussion. REW wrote the manuscript draft with support from JMD. All authors provided constructive critiques in editing the manuscript. All authors contributed to the article and approved the submitted version.

## 7 Funding

REW is supported by 1F31CA261056-01 through NIH National Cancer Institute. HDN is supported by the American Society of Hematology Scholar Award, the National Institutes of Health’s National Center for Advancing Translational Sciences, grants KL2TR002492 and UL1TR002494, a grant from the National Heart, Lung, and Blood Institute under award number R01HL163011, and the 2022 AACR Career Development Award to Further Diversity, Equity, and Inclusion in Cancer Research, which is supported by Merck, Grant Number 22-20-68-NGUY. JMD is supported by the National Cancer Institute of the National Institutes of Health under award number R01CA269801 and the Department of Defense Prostate Cancer Research Program W81XWH-18-1-0542. The content is solely the responsibility of the authors and does not necessarily represent the official views of the National Institutes of Health.

## 8 Acknowledgments

We thank members of the Drake lab for providing advice and input on the manuscript. We also thank Dr. Scott Dehm at the University of Minnesota for his gifts of AD1 and R1D567 cells.

## 9 Supplementary Material

Supplementary Material should be uploaded separately on submission, if there are Supplementary Figures, please include the caption in the same file as the figure. Supplementary Material templates can be found in the Frontiers Word Templates file.

Please see the Supplementary Material section of the Author guidelines for details on the different file types accepted.

## 10 Data Availability Statement

The datasets [GENERATED/ANALYZED] for this study can be found in the [NAME OF REPOSITORY] [LINK]. Please see the “Availability of data” section of Materials and data policies in the Author guidelines for more details.

