## Supplemental Figures for "Saracatinib synergizes with enzalutamide to downregulate androgen receptor activity in castration resistant prostate cancer"

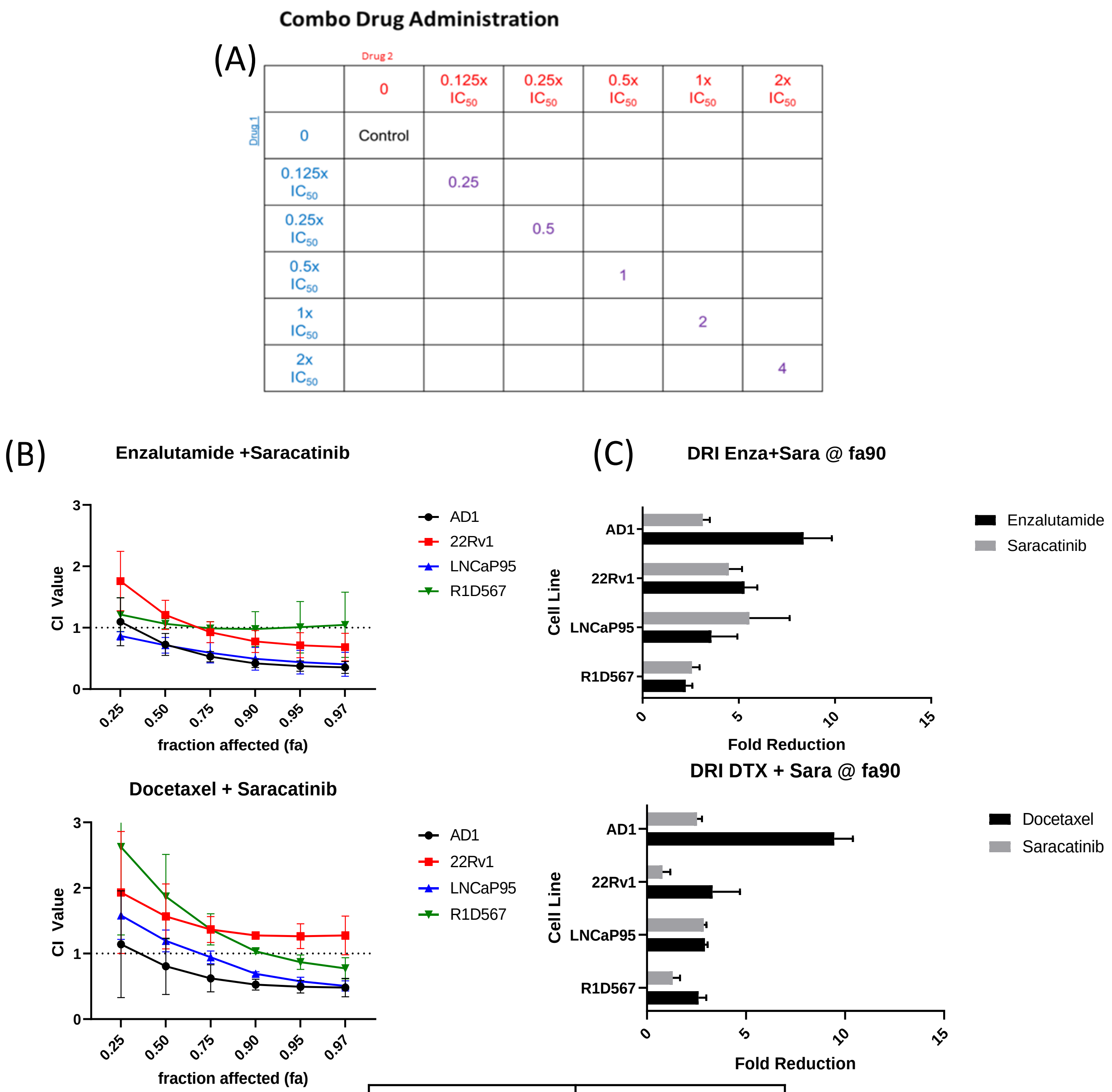


| | |
|---|---|
| CI > 1 | Antagonistic |
| CI = 1 | Additive |
| CI < 1 | Synergistic |

**Supplementary Figure 1: Synergy observed between Enzalutamide and SRC Kinase Inhibitors in AR+ Positive Cell Lines.** (A) Template layout for combination drug administration for cell lines. (B) Combination Index via Chou Talalay results for Enza plus Sara combination and DTX plus Sara combination in AR+ cell lines (AD1, 22Rv1, LNCaP95, R1D567). (C) Dose Reduction Index at 90% of each cell line affected with the enza plus sara combination and the DTX plus sara combination . N≥ 3 for all cell lines except LNCaP95, where N=2.

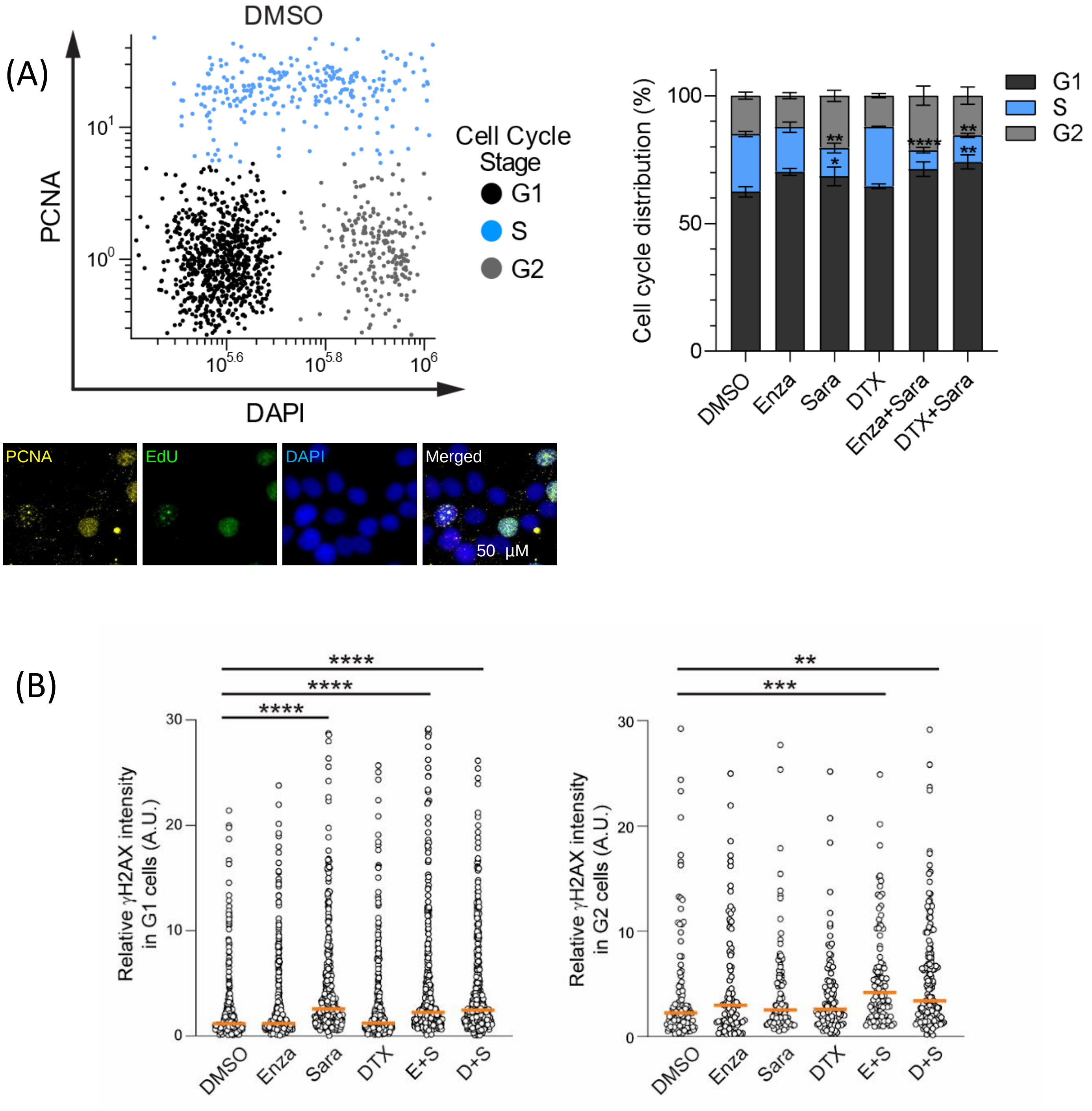


**Supplemental Figure 2: Saracatinib halts DNA synthesis, induces DNA Damage and activates markers of apoptosis a)** Quantitative Image-based cytometry results comparing PCNA to DAPI to identify G1, S-phase, and G2 cells. **b)** γH2AX quantification by G1 and G2. N=3. *, $P < 0.05$; **, $P < 0.01$; ***, $P < 0.001$; and ****, $P < 0.0001$.
